# Precuneus activity during retrieval is positively associated with amyloid burden in cognitively normal older *APOE4* carriers

**DOI:** 10.1101/2024.07.18.604145

**Authors:** Larissa Fischer, Eóin N. Molloy, Alexa Pichet Binette, Niklas Vockert, Jonas Marquardt, Andrea Pacha Pilar, Michael C. Kreissl, Jordana Remz, Judes Poirier, M. Natasha Rajah, Sylvia Villeneuve, PREVENT-AD Research Group, Anne Maass

## Abstract

The precuneus is an early site of amyloid-beta (Aβ) accumulation. Previous cross-sectional studies reported increased precuneus fMRI activity in older adults with mild cognitive deficits or elevated Aβ. However, longitudinal studies in early Alzheimer’s disease (AD) risk stages are lacking and the interaction with Apolipoprotein-E (*APOE*) genotype is unclear. In the PREVENT-AD cohort, we assessed how precuneus activity during successful memory retrieval at baseline and over time relates to future Aβ and tau burden and to change in memory performance. We further studied the moderation by *APOE4* genotype. We included 165 older adults (age: 62.8±4.4 years; 113 female; 66 *APOE4* carriers) who were cognitively normal at baseline and had a family history of AD. All participants performed task-fMRI at baseline and underwent ^18^F-flortaucipir-PET and ^18^F-NAV4694-Aβ-PET on average 5 years later. We found that higher baseline activity and greater longitudinal change in activity in precuneus were associated with higher subsequent Aβ in *APOE*4 carriers but not non-carriers. There were no effects of precuneus activity on tau burden. Finally, *APOE4* non-carriers with low baseline activity in the precuneus exhibited better longitudinal performance in an independent memory test compared to *APOE4* non-carriers with high baseline activity and *APOE4* carriers. Our findings suggest that higher task-related precuneus activity at baseline and over time are associated with subsequent Aβ burden in cognitively normal *APOE4* carriers. Our results further indicate that the absence of hyperactivation and the absence of the *APOE4* allele is related with the best future cognitive outcome in cognitively normal older adults at risk for AD.

**Significance Statement:** The precuneus is a brain region involved in episodic memory function and is an early site of amyloid-beta (Aβ) accumulation. Alterations in task-related activity occur in the precuneus with ageing as well as with Alzheimer’s disease (AD) pathology even in the absence of cognitive symptoms; however, their course and implications are not well understood. We demonstrate that higher precuneus activity at baseline and its change over time during successful memory retrieval is associated with higher Aβ burden on average 5 years after baseline in Apolipoprotein-E4 (*APOE4)* carriers. Lower precuneus baseline activation was related to better memory performance over time in *APOE4* non-carriers. Our findings provide novel longitudinal evidence that increased activity in posterior midline regions is linked to early AD pathology in dependence of *APOE4* genotype.

## Introduction

Changes in brain activity occur naturally throughout the lifecourse. Changes also occur early over the course of Alzheimer’s disease (AD) and can be measured indirectly with functional magnetic resonance imaging (fMRI). Understanding these early changes and how they are mechanistically linked to pathophysiology and progression of AD represents a key goal of cognitive neuroscientists and offers opportunities for preventing cognitive decline (Corriveau-Lecavalier et al., 2024). The posteromedial cortex (PMC) is a key brain region in this regard and includes the precuneus, which itself is among the earliest regions affected by amyloid (Aβ) pathology (Villeneuve et al., 2015; Palmqvist et al., 2017). The precuneus is therefore a promising region to investigate early aberrant activity. Further, it is strongly involved in episodic memory processing (Cavanna and Trimble, 2006; Elman et al., 2013; Moscovitch et al., 2016), a domain that declines both healthy ageing and early in AD (Hedden and Gabrieli, 2004; Rönnlund et al., 2005; McKhann et al., 2011).

Several lines of research suggest that precuneus dysfunction plays a role in cognitive ageing and AD pathogenesis. Previous research has shown greater encoding-related precuneus deactivation in young adults relative to older adults (Lustig et al., 2003), suggesting heightened activation with age. Interestingly, this finding has been replicated in other cohorts (Miller et al., 2008; Pihlajamäki et al., 2008; Bejanin et al., 2012; Mormino et al., 2012; Kizilirmak et al., 2023) and tasks, including an autobiographical memory retrieval task (Fenerci et al., 2022). Similarly, increased precuneus activity during encoding has also been observed in older adults with subjective cognitive decline (SCD) (Corriveau-Lecavalier et al., 2020; Billette et al., 2022), individuals that are at increased risk of developing AD dementia (Mitchell et al., 2014; Slot et al., 2019). Several studies in cognitively normal older adults reported associations between reduced precuneus deactivation during encoding and higher Aβ burden (Sperling et al., 2009; Vannini et al., 2012). Further studies have also found associations of the Apolipoprotein-E4 (*APOE4)* genotype (Han et al., 2007; Persson et al., 2008; Pihlajamäki et al., 2010) and heightened activity. *APOE4* is a major risk factor for AD (Mayeux, 2003; Liu et al., 2013) and strongly correlated with Aβ accumulation (Villemagne and Rowe, 2013; Selkoe and Hardy, 2016). Recently, it has been proposed that *APOE4* homozygosity represents a distinct form of genetic AD, with almost all homozygotes showing AD pathology and cognitive symptoms in their later life (Fortea et al., 2024). Therefore, there seems to be accumulating evidence for a role of aberrant hyperactivation of PMC regions, in addition to the well-established risk associated with *APOE4* genotype, in the preclinical stages of AD. However, how these two factors interact to affect the spread of AD pathology remains unclear and to be empirically tested.

Here we assessed the relationship between precuneus fMRI retrieval activation at baseline and change over time, *APOE4* genotype, cognitive changes, and the level of AD pathology in cognitively normal older adults from the longitudinal Pre-symptomatic Evaluation of Experimental or Novel Treatments for Alzheimer’s Disease (PREVENT-AD) cohort. PREVENT-AD incorporates multimodal data from cognitively unimpaired older adults with a familial history of sporadic AD (Tremblay-Mercier et al., 2021). We hypothesised that i) precuneus brain activity would be higher or increasing over time in *APOE4* carriers compared to non-carriers, ii) increased precuneus activity at baseline and increases over time would be linked to future Aβ and tau burden, iii) higher activity or activity changes would be positively (if beneficial) or negatively (if detrimental) linked to cognitive changes, and iv) *APOE* genotype might moderate associations between activity, pathology, and cognition.

## Methods

### Sample and study design

All participants were cognitively unimpaired older adults from the open science PREVENT-AD cohort study launched in 2011 (Breitner et al., 2016; Tremblay-Mercier et al., 2021). They had at least 1 parent or 2 siblings diagnosed with AD-like dementia, which is associated with an increased risk for developing sporadic AD (Donix et al., 2012). Participants were above 60 years of age at baseline. People aged between 55 and 59 were also included if they were less than 15 years away from the age of onset of symptoms of their first-affected relative. Participants had no major neurological or psychiatric illnesses at time of enrolment. Inclusion criteria comprised intact cognition based on the Montreal Cognitive Assessment (MoCA) questionnaire with a score of at least 26 of 30 points (Nasreddine et al., 2005), a Clinical Dementia Rating (CDR) Scale of 0 (Morris, 1993) or exhaustive neuropsychological evaluation. All participants included in our analyses underwent a baseline fMRI scan, cognitive assessments using the standardised Repeatable Battery for the Assessment of Neuropsychological Status (RBANS) (Randolph et al., 1998) and both an Aβ and tau PET scan at varying times (mean=5 years, range=0.5-10 years) post-baseline fMRI scan. This selection created a subsample of 165 participants (aged 62.8±4.4 years at baseline, 15.42±3.3 years of education, 52 male/ 113 female, 66 *APOE4* carriers including 3 *APOE4* homozygotes) upon which our analyses were performed. Follow-up fMRI scans and RBANS assessments were performed over the course of 48-months in a subset of participants. Specifically, participants underwent a 3-month (N=79), 12-month (N=135), 24-month (N=111), and 48-month (N=58) follow-up fMRI scan after baseline (Fig.1). All study procedures and experimental protocols were approved by the McGill University Institutional Review Board and/ or the Douglas Mental Health University Institute Research Ethics Board. All participants provided written informed consent prior to each experimental procedure and were financially compensated for their time.

**Fig. 1.**
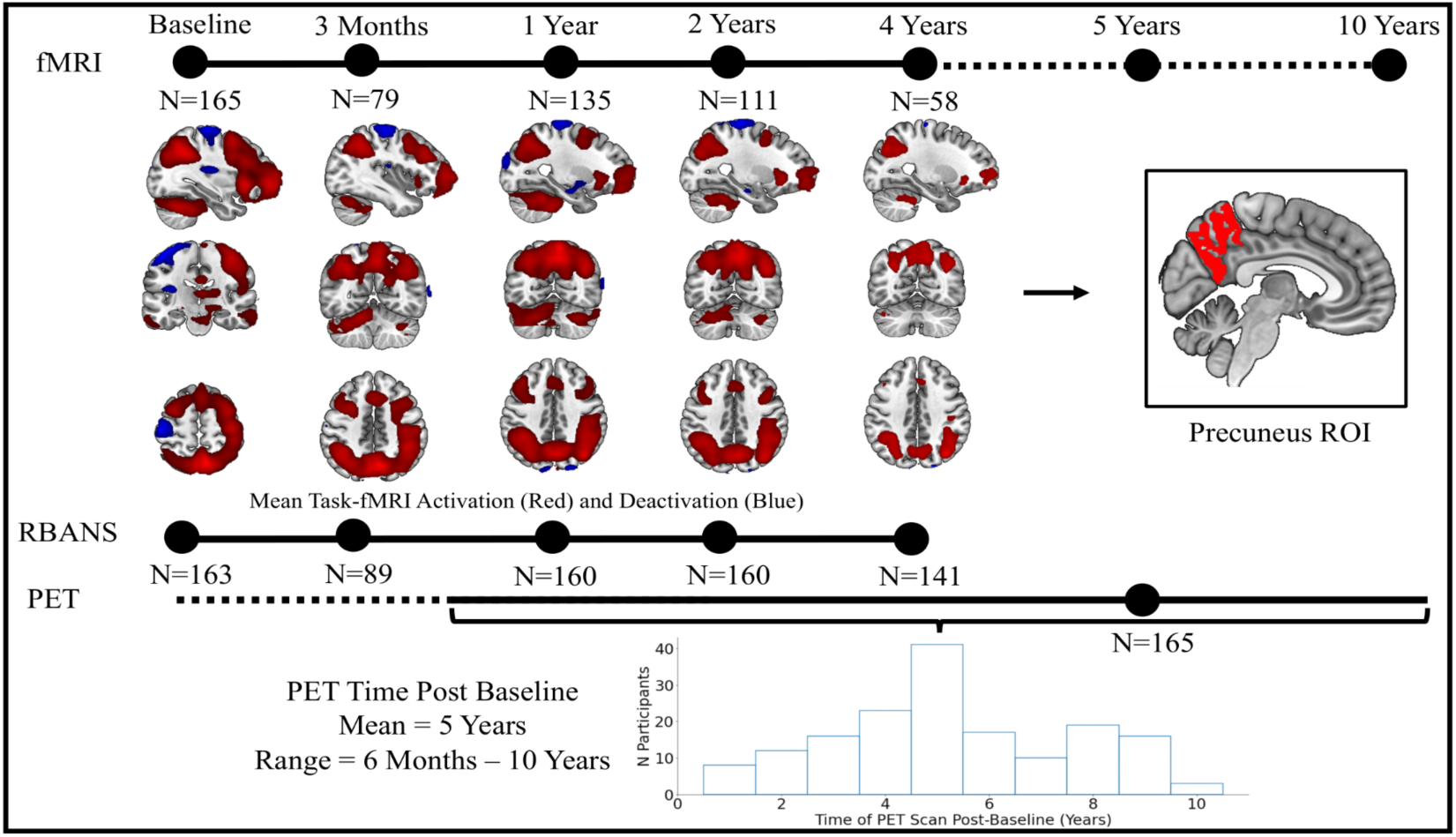
Study Design: Each participant underwent one baseline fMRI session and up to 4 fMRI follow-up sessions with the last follow-up 4 years after baseline. Similarly, RBANS neuropsychological assessments were performed at baseline and over time. All 165 participants underwent PET scans to quantify amyloid-beta and tau pathology between 6 months and up to 10 years after the baseline fMRI scan. Group fMRI activity during successful retrieval (red scale: hits > correct rejections; inverse contrast in blue) is depicted for each time-point. FMRI results shown at *p*<0.05 FWE-corrected at the voxel level. fMRI = Functional Magnetic Resonance Imaging, RBANS = Repeatable Battery for the Assessment of Neuropsychological Status, PET = Positron Emission Tomography.

### Task-fMRI design

FMRI data were acquired using a Siemens Tim Trio 3-Tesla MRI scanner at the Cerebral Imaging Centre of the Douglas Mental Health University Institute using a Siemens standard 12 or 32-channel coil (Siemens Medical Solutions, Erlangen, Germany) (Tremblay-Mercier et al., 2021). Scans were acquired with a TR=2000 ms; TE=30 ms; 90° flip angle, FOV=256x256 mm field of view covering 32 slices, and a 4 mm isotropic voxel resolution. Participants performed an encoding and retrieval block of an object-location episodic memory task within each scan session. Details of the task fMRI methods have been previously published (Rabipour et al., 2020; Tremblay-Mercier et al., 2021). During the encoding task, participants were presented with 48 visual stimuli (coloured line-drawings of everyday items), presented on either the right or left side of the screen. Participants were asked to indicate on which side of the screen the stimulus was presented by pressing a button. After a 20-minute interval of structural scanning, participants performed the retrieval task. They were presented with 48 old stimuli, i.e. object stimuli shown during the encoding session, and 48 new object stimuli. Specifically, participants were asked to indicate via a button press (forced-choice between four alternative answers), whether i) “The object is FAMILIAR but you don’t remember the location” (“F”); ii) “You remember the object and it was previously on the LEFT” (“L”); iii) “You remember the object and it was previously on the RIGHT” (“R”); and iv) “The object is NEW” (“N”) (see supplementary Fig.S1). The retrieval task took approximately 15 minutes. For the purpose of this paper, we focus on brain activity associated with successful object recognition, that is activity differences between correctly recognized (old) objects (irrespective of location/ source memory) versus activity during correct rejection of novel objects.

### Task-fMRI preprocessing and data preparation

All data were preprocessed using MATLAB and Statistical Parametric Mapping, version 12 (SPM12) (Functional Imaging Laboratory UCL, n.d.). Data were realigned, slice time-corrected, co-registered to an anatomical T1 image, normalised, and smoothed using a 8 mm full-width-half-maximum (FWHM) Gaussian kernel. Three-dimensional T1 anatomical data (TR=2300 ms; TE=2.98 ms; TI=900 ms; 9° flip angle; FOV=256x240x176 mm) with a 1 mm isotropic voxel resolution were segmented for functional image normalisation using the unified segmentation approach (Ashburner and Friston, 2005). Following preprocessing, we performed first-level analyses. The first-level GLM included 3 regressors of interest: hits (responses “F”/”L”/”R” to old object stimuli), correct rejections (response “N” to new object stimuli) and false alarms or misses (all other responses) as well as six motion regressors from the realignment process. All included participants had a minimum of 10 hits and 10 correct rejections, specifying a *t*-contrast, hereafter referred to as the episodic memory contrast. To specify the episodic memory contrast, we compared “hits” (previously viewed items that were correctly identified, regardless of their previously presented location on screen) with “correct rejections” (new items correctly identified as new). To assess precuneus brain activity associated with the episodic memory contrast, we applied a region of interest (ROI) approach using FreeSurfer (Laboratories for Computational Neuroimaging, Athinoula A. Martinos Center for Biomedical Imaging, n.d.) masks (Fig.1) (labels 1025 and 2025 from the aparc+aseg.nii in MNI space), resliced to match functional image dimensions using the “Coregister and Reslice” command in SPM12. Using these masks, we subsequently extracted mean beta values for the bilateral precuneus during the Hits > Correct Rejections episodic memory contrast for each participant using in house MATLAB scripts.

### PET acquisition and preprocessing

PET scans were performed at the McConnell Brain Imaging Centre of the Montreal Neurological Institute (Quebec, Canada) using a brain-dedicated PET Siemens/CTI high-resolution research tomograph. Data acquisition and processing was carried out as previously described (Yakoub et al., 2023). In brief, Aβ-PET images using ^18^F-NAV4694 (NAV) were acquired 40 to 70 min after injection, with an injection dose of ∼6 mCi. Tau-PET images, using ^18^F-flortaucipir (FTP), were acquired 80 to 100 min after injection, with an injection dose of ∼10 mCi. Frames of 5 minutes as well as an attenuation scan were obtained. PET images were reconstructed using a 3D ordinary Poisson ordered subset expectation maximum algorithm (OP-OSEM), with 10 iterations, 16 subsets, while all images were decay and motion corrected. Scatter correction was performed using a 3D scatter estimation method. T1-weighted MRI images were parcellated into 34 bilateral ROIs based on the Desikan-Killiany atlas using FreeSurfer version 5.3. PET images were realigned, temporally averaged, and co-registered to the T1-weighted image (using the scan closest in time to PET data acquisition), then masked to remove signal from cerebrospinal fluid (CSF) and smoothed with a 6 mm^3^ Gaussian kernel. Standardised uptake value ratios (SUVR) were computed as the ratio of tracer uptake in the ROIs versus uptake in cerebellar grey matter for amyloid-PET scans or versus inferior cerebellar grey for tau-PET. All PET data were preprocessed using a standard pipeline (https://github.com/villeneuvelab/vlpp). We focused on a ROI approach for tau, assessing bilateral entorhinal FTP SUVR, obtained by averaging the uptake ratio of both the left and right entorhinal cortices and whole-brain NAV SUVR.

### *APOE* genotyping

All participants were genotyped for *APOE* using a QIASymphony apparatus, as described previously (Tremblay-Mercier et al., 2021). If participants showed at least one copy of the *APOE4* risk allele, they were allocated to the carrier group while those without were allocated to the non-carrier group (carriers=66 with 22 male, non-carriers=99 with 30 male).

### Assessment of memory performance

We focused on two measures of episodic memory, the RBANS delayed memory index score and corrected hit rate derived from the fMRI retrieval task. The RBANS delayed memory index score is a combined measure of word-list recognition and delayed figure, story and word-list recall (Randolph et al., 1998). Corrected hit rate was specified as hits (i.e. responses “Familiar”, ”Remember-Left” or ”Remember-Right” to previously shown objects) minus false alarms (responses “Familiar”, ”Remember-Left” or ”Remember-Right” to novel objects) during the fMRI retrieval recognition task. We note that different versions of the RBANS were used in follow-up sessions to reduce practice effects and different object stimuli were employed at each fMRI visit. RBANS data from baseline (N=163), a 3-month (N=89), 12-month (N=160), 24-month (N=160), and 48-month (N=141) follow-up were included.

### Statistical analysis

All statistical analyses were conducted using R (R Core Team, 2022), version 4.1.2, implemented within RStudio (RStudio Team, 2022), and running on MAC OS Monterrey version 12.4. The R code used for analyses is publicly available (https://github.com/fislarissa/precuneus_retrieval_hyperactivation). For the linear models, we ensured that heteroscedasticity and multicollinearity were not present. Furthermore, we tested for a normal distribution of residuals using the Shapiro-Wilk test on the standardised residuals of each model. Our analyses focused on 4 major questions: i) Are there differences in precuneus activity during memory retrieval at baseline or over time between *APOE4* carriers and non-carriers?; ii) Is higher precuneus activity at baseline and increase in activity over time associated with future amyloid and tau burden?; iii) Is higher activity or longitudinal activity change positively (if beneficial) or negatively (if detrimental) related to cognitive changes?; and iv) Does *APOE* genotype moderate any association between activity and pathology or cognition (e.g. is higher activity related to more pathology only in *APOE4* carriers)?

### Assessment of baseline and longitudinal precuneus activation and *APOE* genotype

Following extraction of precuneus-specific magnitude of brain activity, we specified a linear model to assess effects of *APOE4* status adjusting for age, sex, precuneus grey matter volume (GMV) at baseline, obtained from T1-weighted structural images, and years of education. We then specified a linear mixed model (LMM) (Bates et al., 2015) to investigate changes in activity over time, with precuneus activity as the dependent variable and time (as the scaled continuous time difference between individual sessions) as the fixed within-subject effect and random intercepts per participant, adjusting for the same variables. We then repeated the analysis including an interaction term of time by *APOE* status and as a supplementary analysis time by sex.

### Assessment of baseline and longitudinal precuneus activity, AD pathology, and *APOE* genotype

To test whether baseline precuneus activity predicted AD pathology at follow-up, we specified two linear models in which baseline precuneus activity was used as the independent and (i) whole-brain amyloid PET and (ii) entorhinal tau SUVRs as the dependent variables, respectively. Age at baseline, sex, years of education, time (in months) between the baseline fMRI scan and the respective PET scan, and precuneus GMV at baseline were specified as covariates in each model. Due to the non-normal distribution of amyloid and tau pathology, we applied a Box-Cox transformation to the PET data in order to achieve a closer approximation of a normal distribution. Following the analyses of the effects of baseline activity on AD pathology, we extracted the slope of the change in precuneus activity over time for each participant. The specified model for the slope extraction included precuneus activation as dependent variable, time (as the scaled continuous time difference between individual sessions) as independent variable, and a random intercept and slope per participant. Subsequently, we entered the extracted slope of activation as the predictive variable in a second set of linear regressions, assessing the effects of change in precuneus activity over time on AD pathology at follow-up. Age, sex, education, and precuneus GMV at baseline were again included as covariates. To assess whether the activity slope was associated with the baseline fMRI signal, we performed a correlation analysis.

In a subsequent analysis considering *APOE* genotype, we first tested for a difference in amyloid and tau burden between *APOE* genotype groups, adjusting for age, sex and education. We subsequently repeated our linear regression analyses in which we tested if there was an interaction between *APOE4* genotype and precuneus activation at (i) baseline and (ii) over time (slope) on AD pathology at follow-up.

### Assessment of baseline precuneus activation and baseline episodic memory retrieval performance as well as longitudinal episodic memory retrieval performance

To test for associations between baseline precuneus activation and baseline corrected hit rate (specified as hits minus false alarms) of fMRI task-performance and the delayed memory score recorded on the RBANS, we used partial correlation analyses (correcting for years of education, sex and age). We then modelled the longitudinal corrected hit rate from the task fMRI or RBANS delayed episodic memory as the dependent variable in two LMM and time as the within-subject factor. Age, sex, and years of education were covariates in all analyses.

### Assessment of baseline precuneus activation, *APOE* genotype, and longitudinal episodic memory retrieval performance

In order to assess the interaction effects of genotype and precuneus activation on cognitive measures of episodic memory over time, we created two LMMs in which episodic memory performance (first measured by the corrected hit rate from the task fMRI and second from the RBANS delayed memory) was specified as the dependent variable, precuneus activation at baseline as the independent variable and session (as session 1 to 5; to investigate session-specific differences) and APOE genotype were specified as within-subject factors. Again, this model was specified with age, sex, and education as covariates. We first investigated the interaction effects of APOE genotype by session and baseline activity by session, then a three-way-interaction of APOE genotype by baseline activity by session on memory performance. We then applied post hoc contrasts to each session for those models with significant interactions. Finally, we investigated group effects on the performance slopes over time (as the scaled continuous time difference between individual sessions) correcting for age, sex and education, specifically APOE4 carriers vs. non-carriers for the RBANS delayed memory score and APOE4 non-carriers with low baseline activation vs. all other groups for corrected hit rate performance.

## Results

### Assessment of baseline and longitudinal precuneus activation and *APOE* genotype

Precuneus activity during memory retrieval at baseline in all (N=165) participants did not differ due to *APOE4* status (*p*>0.05; see supplementary Tab.S1). Regarding longitudinal changes in precuneus activity, there was a statistically significant decrease of precuneus retrieval activity over time (β=-0.15 [95% CI -0.22, -0.07], t=-3.987, *p*<0.001) in all participants with >1 fMRI scan (N=151). There was also no significant time by *APOE* group interaction (*p*>0.05; see supplementary Tab.S2), with both carriers and non-carriers exhibiting similar decreases in precuneus brain activity over time (Fig.2A). For significant sex effects, see supplementary Fig. S2.

**Fig. 2.**
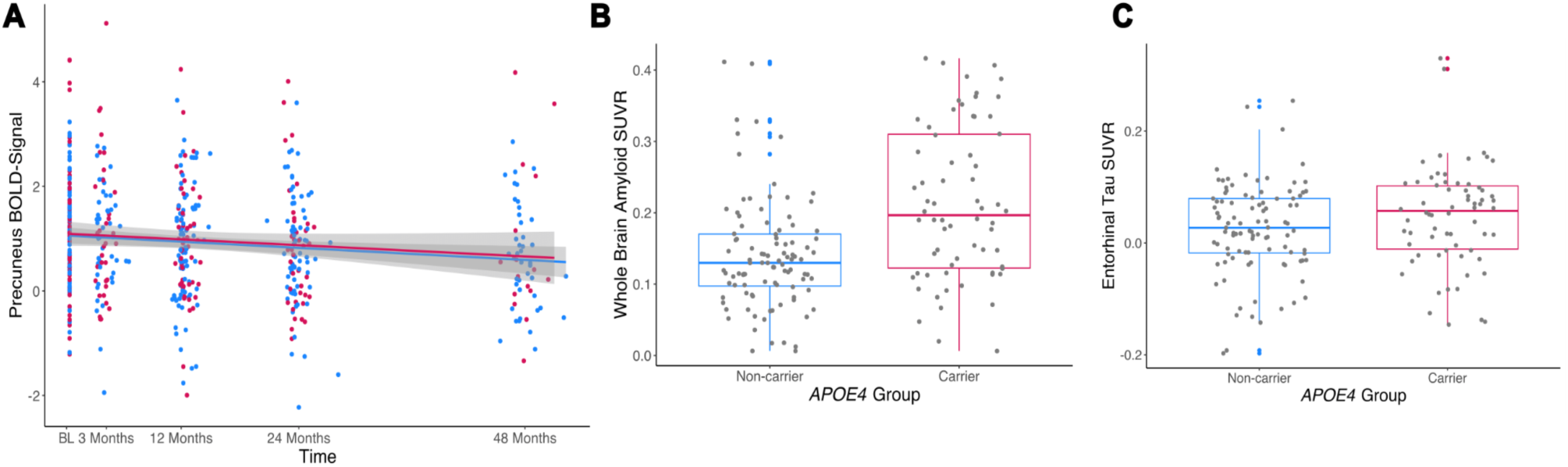
Precuneus retrieval activity over time accounting for *APOE* genotype and Alzheimer’s pathology differences between *APOE* genotype groups: **A**: Linear mixed modelling showed a significant decrease in precuneus activity over time in the whole sample. There was no significant time by *APOE* genotype interactions, suggesting comparable changes over time in *APOE4* carriers (red) and non-carrier (blue). Shaded areas refer to the 95% confidence interval. **B:** A group comparison of whole-brain amyloid burden showed a significantly higher burden for *APOE4* carriers versus non-carriers, when adjusting for age, sex, and years of education. **C:** Similarly, *APOE4* carriers also showed a marginally higher tau burden in the entorhinal cortex. BL = Baseline.

### Assessment of baseline and longitudinal precuneus activation, AD pathology, and *APOE* genotype

Linear regression analyses assessed the predictive effects of precuneus baseline activation as well as change of activation (slope over time derived from LMM) on PET measures of AD pathology on average 5 years after baseline. With regard to Aβ burden, we found that higher baseline precuneus activity was related to significantly higher whole-brain NAV SUVR (β=0.20 [95% CI 0.05, 0.36], t=2.544, *p*=0.012; Fig.3A) in the whole sample. A similar model with activity change as predictor yielded a significant positive association between precuneus activity slope and PET-measured whole-brain Aβ (β=0.17, [95% CI 0.01, 0.34], t=2.082, *p*=0.039; Fig.3B). The analyses assessing the relationship between precuneus activation and bilateral entorhinal tau, however, did not yield a significant result for the baseline precuneus activation (β=0.08, [95% CI -0.08, – 0.23], t=0.972, *p*=0.332). Similarly, we did not observe a significant predictive effect of precuneus slope on bilateral entorhinal tau (β=0.08, [95% CI -0.09, 0.25], t=0.954, *p*=0.342). Additionally, baseline precuneus retrieval activation and the slope of activation over time were positively correlated (r=0.73, [95% CI 0.64, 0.79, t(149)=12.913, *p*<0.001). Regarding AD pathology burden, *APOE* carriers exhibited significantly higher whole-brain Aβ (β=0.79, [95% CI 0.50, 1.09], t=5.335, *p*<0.001; Fig.2B) and entorhinal tau burden (β=0.32, [95% CI 0.01, 0.63], t=2.040, *p*=0.043; Fig.2C).

**Fig. 3.**
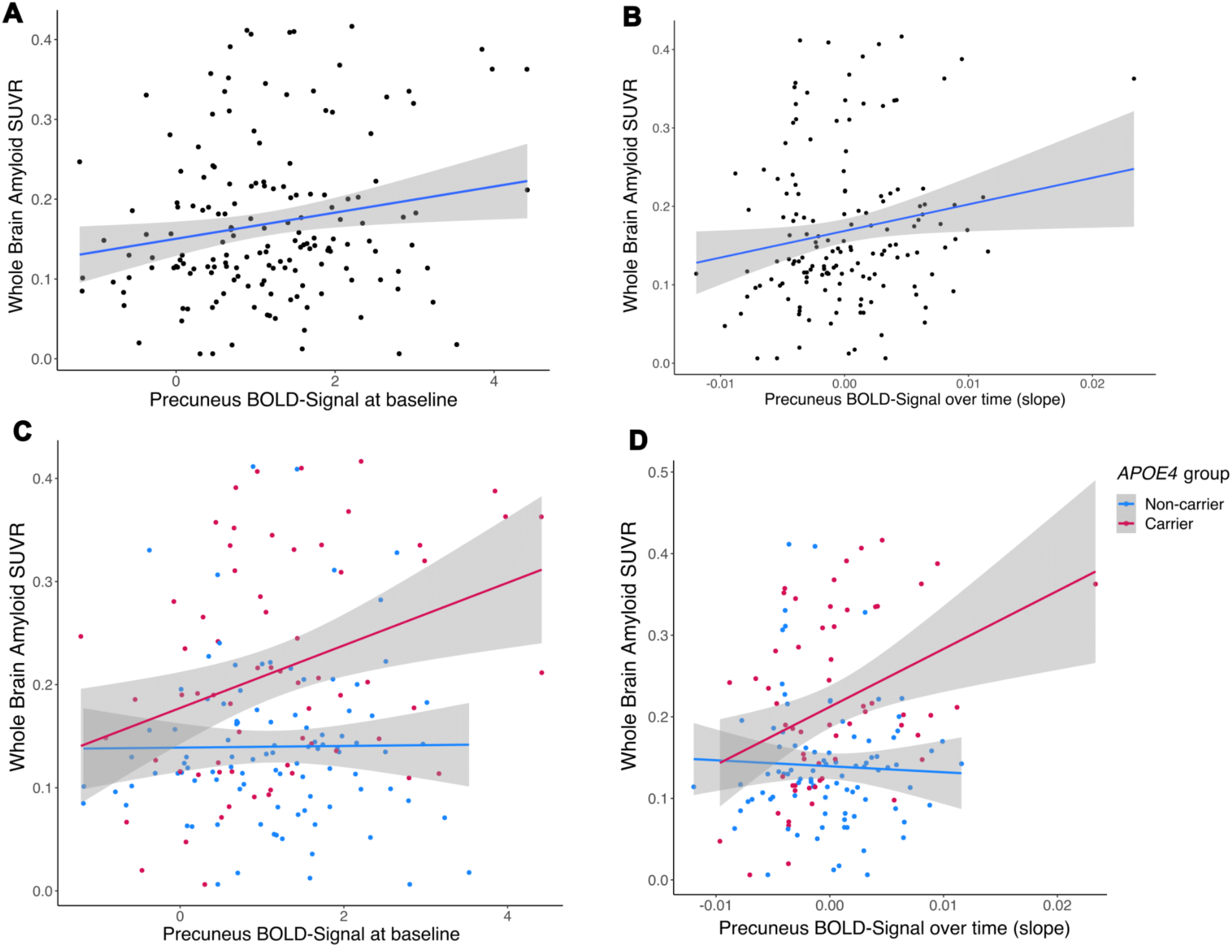
Relationship between precuneus activation and PET-assessed measures of amyloid burden. **A:** A linear regression model showed that baseline precuneus activation was significantly related to later whole-brain amyloid burden. **B:** Change in precuneus activation over time was also significantly associated with amyloid PET burden, with higher slope values being associated with higher amyloid burden. **C:** *APOE* genotype moderated the association between precuneus activation at baseline and future whole brain amyloid burden (with higher brain activation levels at baseline predicting higher levels of amyloid in *APOE4* carriers). **D:** Similarly, an interaction between APOE4 genotype and precuneus activation over time (slope) was observed, with weaker decreases or increases in precuneus activation being associated with higher subsequent amyloid burden in *APOE4* carriers (red dots). Shaded areas refer to the 95% confidence interval.

Next, we assessed the potential moderating effect of *APOE* genotype on the relationship between precuneus baseline activity or activity change and AD burden, thereby including genotype as a group factor in the model. Regarding Aβ burden, the interaction between baseline precuneus activity and *APOE* genotype on whole-brain NAV SUVR was significant (β=0.29, [95% CI 0, 0.57], t=2.004, *p*=0.047; Fig.3C), such that *APOE4* carriers with higher baseline activation showed higher future Aβ-PET burden (β=0.33, [95% CI 0.08, 0.58], t=2.622, *p*=0.011; Fig.3C, red line). Similarly, there was a significant precuneus activity slope by *APOE* genotype group interaction (β=0.39, [95% CI 0.10, 0.69], t=2.631, *p*=0.009) on whole-brain NAV SUVR (Fig.3D), meaning *APOE4* carriers with weaker activity decreases (or more activity increases) over time showed higher subsequent Aβ-PET burden (β=0.36, [95% CI 0.10, 0.63], t=2.758, *p*=0.008; Fig.3D, red line). In contrast, baseline precuneus activity or longitudinal change in precuneus activity were not related to future Aβ-PET burden in non-carriers (*p*>0.05; Fig.3C and D, blue lines). Statistical details are provided in supplementary Tab.S3-6. With respect to entorhinal tau burden, no significant interaction effects between *APOE* genotype and precuneus activity, neither baseline nor slope (all *p*>0.05), on FTP SUVR were observed.

### Assessment of baseline precuneus activation and baseline episodic memory retrieval performance as well as longitudinal episodic memory retrieval performance

Partial correlations showed no significant association of baseline precuneus task-activation with baseline task-fMRI episodic memory performance (rho=0.13, *p*=0.09) or with the baseline RBANS delayed memory index score (rho=-0.01, *p*=0.80). FMRI task performance as measured by corrected hit rate did not change significantly over time (β=-0.06, [95% CI -0.13, -0.01], t=-1.604, *p*=0.109). However, there was a significant effect of sex (β=-0.33, [95% CI -0.58, -0-08], t=-2.597, *p*=0.010) with males showing overall worse performance. No effect of age or education was present (*p*>0.05). Similar modelling of cognitive performance measured by the RBANS delayed memory index score showed a significant increase over time (β=0.11, [95% CI 0.06, 0.17], t=3.915, *p*<0.001). Additionally, we observed a significant effect of sex (β=-0.44, [95% CI -0.67, -0.21], t=-3.771, *p*<0.001) and education (β=0.22, [95% CI 0.11, 0.32], t=4.143, *p*<0.001) on longitudinal RBANS delayed memory performance, such that male participants showed lesser improvements in performance over time than females and higher education predicted greater performance increases. No effect of age was observed (*p*>0.05).

### Assessment of baseline precuneus activation, *APOE* genotype, and longitudinal episodic memory retrieval performance

We assessed whether baseline precuneus activity, *APOE* genotype or their interaction predicted longitudinal change in memory performance, which showed different results for memory performance for the corrected hit rate and the RBANS delayed memory index. In an LMM that included an *APOE* genotype by session and a baseline precuneus activity by session interaction on memory performance, there was a significant *APOE* genotype by session interaction on corrected hit rate (F(4, 435)=2.679, *p*=0.031; see supplementary Fig.S3), but not on RBANS (*p* >0.05). Regarding the slopes over time for the corrected hit rate, there was a significant effect of *APOE* group (F(1,148)=89.537, *p*<0.001), with *APOE4* carriers (mean=-0.03, SD=0.02) showing a steeper slope (i.e. decline over time) than non-carriers (mean=-0.01, SD=0.01) as shown in supplementary Fig.S3. Post-hoc analyses per session revealed that *APOE4* carriers had a higher corrected hit rate compared to non-carriers at the 3-month follow-up session (t(546)=-2.326, *p*=0.020, SE=0.05, [95% CI -0.20, -0.02]). However, non-carriers had statistically non-significant higher performance than APOE4 carriers at the 24-month (t(499)=0.856, *p*=0.392, SE=0.04, [95% CI -0.05, 0.12]) and 48-month (t(582)=1.273, *p*=0.204, SE=0.06, [95% CI -0.04, 0.19]) session. There was no baseline activity by session interaction on corrected hit rate or on RBANS (all *p* >0.05), no significant main effects of *APOE* genotype or baseline activity were found, neither for the corrected hit rate nor for the RBANS (all *p* >0.05)

Finally, we tested whether baseline precuneus activation predicted change in memory performance in dependence on *APOE* genotype by extending the previous model by a 3-way interaction (activity by *APOE* genotype by session) There was no three-way interaction for corrected hit rate (*p*>0.05). Regarding the RBANS delayed memory index score, the LMM showed a significant *APOE* genotype by baseline precuneus activation by session interaction (F(4, 561)=2.5852, *p*=0.036; Fig.4) on delayed memory performance. When splitting the two *APOE* groups each into a high- and a low-activation group regarding baseline precuneus activation depending on the value being above or below the mean, *APOE4* non-carriers (Fig.4, blue lines) with lower precuneus activity showed a steeper slope for RBANS over time (corresponding to greater increases in delayed memory; N=50, mean=1.32, SD=0.3) in contrast to all other combinations pooled together. This comprises *APOE4* non-carriers with high baseline activity (N=49, mean=0.78, SD=0.45) and *APOE4* carriers (Fig. 4, red lines) with low (N=37, mean=0.52, SD=0.97) or high baseline activity (N=27, mean=0.63, SD=0.5) (t(156)=6.483, p<0.001, SE=0.31, [95% CI 1.99, 2.60]). Post-hoc analyses on cross-sectional RBANS performance at follow-up visits between *APOE4* groups and high and low baseline precuneus activation (fixed at the 25% and 75% percentile) revealed an interaction of *APOE4* group and session for lower activation (see supplementary Fig.S4). With lower baseline activation, there was higher performance for *APOE4* carriers compared to non-carriers at the 3-month follow up (t(690)=-2.124, *p*=0.034, SE=2.04, [95% CI -8.33, -0.33]; not corrected for multiple comparisons), but descriptively higher performance for non-carriers at the 24-month (t(507)=1.109, *p*=0.268, SE=1.60, [95% CI -1.37, 4.92]) and 48-month (t(544)=1.484, *p*=0.138, SE=1.67, [95% CI -0.80, 5.74]) follow ups. There were no differences in performance between *APOE* groups with higher activation (all p>0.05; see supplementary Tab.S7).

**Fig. 4.**
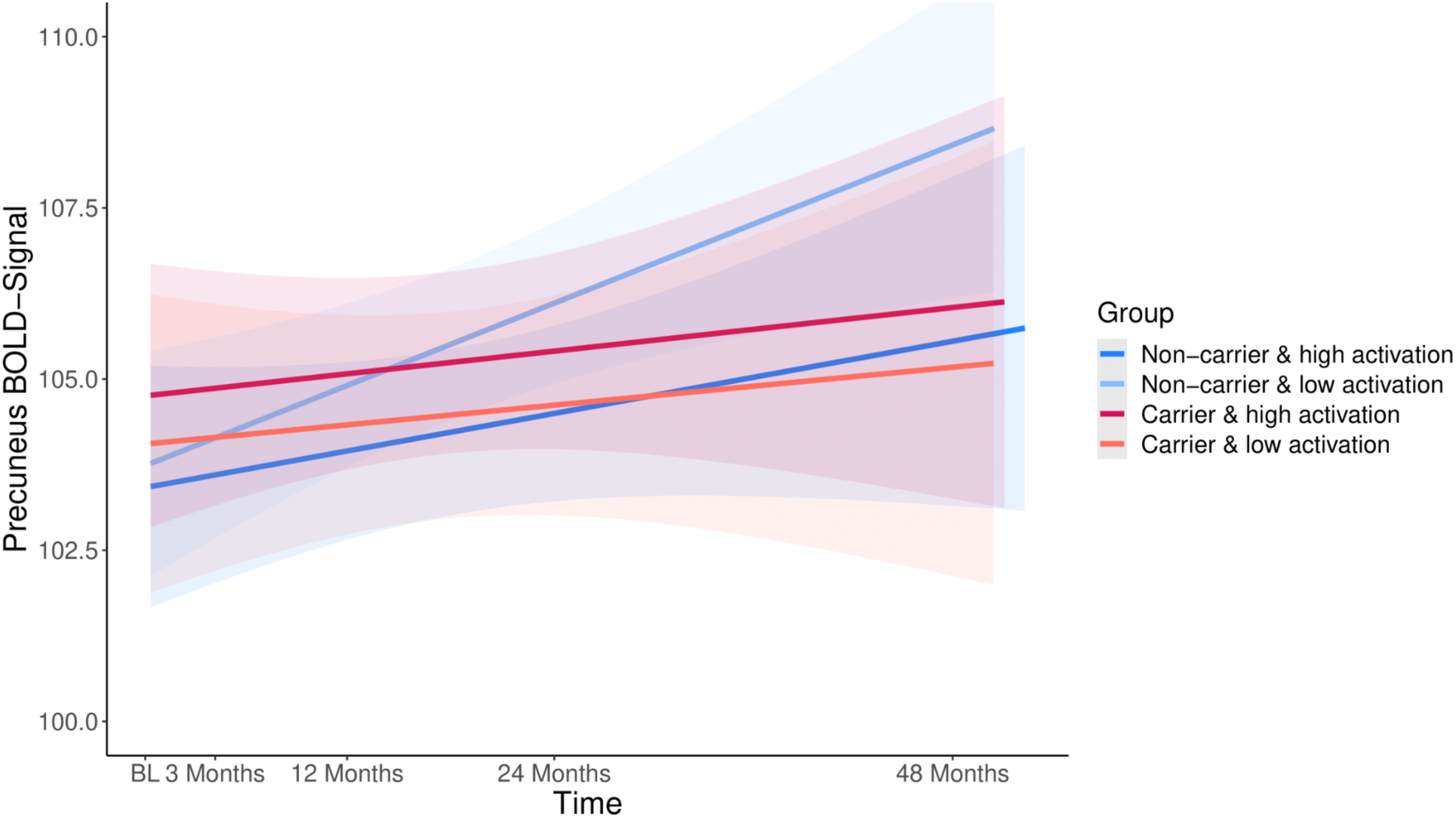
Slope of RBANS delayed memory performance over time considering *APOE* genotype and precuneus baseline activation. *APOE4* non-carriers with lower precuneus activation showed stronger increases (steeper slope) in RBANS performance over time (corresponding to better delayed memory) in contrast to all other combinations (*APOE4* non-carriers with high baseline activation and *APOE4* carriers with low or high baseline activation), suggesting that the absence of the *APOE4* allele and low precuneus activation at baseline are related to the best cognitive outcomes (trajectory). We split the two *APOE* groups each into a high-and a low-activation group regarding baseline precuneus activation depending on the value being above or below the mean. Shaded areas refer to the 95% confidence interval. RBANS = Repeatable Battery for the Assessment of Neuropsychological Status.

## Discussion

In this study, we utilised the longitudinal open science PREVENT-AD dataset to test whether higher precuneus activity at baseline or longitudinal change differed by *APOE4* genotype and whether it was associated with future Aβ burden in cognitively normal older adults. While *APOE4* carriers did not show higher precuneus activity during retrieval per se, higher baseline activation and its change over time were associated with later whole-brain amyloid in this group. We did not, however, observe any effect of precuneus activity or *APOE4* on entorhinal tau pathology, suggesting a specific early predictive effect of activation and *APOE4* for amyloid burden. Finally, we also show a link between brain activity, genotype, and cognitive trajectories, such that *APOE4* non-carriers with low precuneus brain activity at baseline show the strongest improvement or practice effects over time in an independent delayed memory test.

These longitudinal results indicate that increased task-based precuneus activation is associated with a subsequent increase in Aβ burden. Previous cross-sectional studies have reported associations between higher activation (or reduced deactivation) of the PMC during different cognitive tasks and higher amyloid burden (Sperling et al., 2009; Vannini et al., 2012; Elman et al., 2014; Oh et al., 2015), similar to our longitudinal fMRI findings within this cohort. Interestingly, most studies assessed memory encoding activity (for review, see (McDonough et al., 2020; Corriveau-Lecavalier et al., 2024), whereas we investigated increased activity during successful memory retrieval. Mechanistically, animal models suggest that neuronal hyperexcitability, which may translate to aberrantly higher cerebral activation, facilitates amyloid accumulation (Bero et al., 2011) and is also induced by amyloid-related processes (Zott et al., 2019), therefore potentially forming a vicious cycle. This could suggest that the increased BOLD signal measured in human fMRI studies during memory encoding or retrieval might represent neuronal hyperexcitability that is linked to subsequent amyloid accumulation, as shown in animal models. Interestingly, very early amyloid burden has also been reported in the precuneus (Chételat et al., 2013; Villeneuve et al., 2015; Palmqvist et al., 2017), which is a highly connected and metabolically active hub region of the default-mode network (DMN) (Buckner et al., 2008). Dynamic causal modelling approaches suggest that increased task-activation within PMC-regions due to higher amyloid load can drive hyperactivation and tau spread in the medial temporal lobe (MTL) (Giorgio et al., 2023), thereby contributing to detrimental processes. This emphasises the close link between high network activity or connectivity and vulnerability to protein aggregation. While our results support previous findings regarding amyloid, we did not observe associations between fMRI activation in the precuneus and later tau accumulation. While we did not explicitly assess MTL activity, which might be more closely linked to tau, our results suggest a specific mechanism linking hyperactivation in precuneus with later Aβ.

Our results also show an interaction between precuneus activity and *APOE* genotype. Specifically, we observed that *APOE4* carriers with higher baseline and longitudinal precuneus activation exhibited subsequent increases in amyloid burden. Moreover, higher precuneus activity did not relate to amyloid in the absence of the *APOE4* allele. While the specific role of *APOE4* in amyloid accumulation and spread is not fully understood, there is converging evidence for a critical role in various dysfunctional mechanisms that could precipitate AD pathology (Papenberg et al., 2015; Hersi et al., 2017; Najm et al., 2019). Animal models point towards a loss of inhibition in the MTL of *APOE4* carriers that could drive hyperactivation, however not much is known about the precuneus (Nuriel et al., 2017; Najm et al., 2019). While causality remains unclear, our results regarding heightened precuneus activation are congruent with previous findings suggesting aberrant hyperexcitability as an early risk factor for amyloid accumulation. Moreover, our findings stress the moderating role of the *APOE* genotype on the link between increased activity and amyloid pathology, whereby individuals with increased precuneus activity that also carry the *APOE4* allele seem to be particularly vulnerable to amyloid accumulation. Moreover, *APOE4* carriers had higher amyloid burden, thereby replicating previous findings (Chételat and Fouquet, 2013; Liu et al., 2013; Martens et al., 2022). As our *APOE4* carrier group mainly consisted of heterozygotes with only one copy of the *APOE4* allele and only 3 homozygotes, we did not further distinguish these groups.

Another possibility that warrants consideration is increased task-related activity as a mechanism of neuronal or network compensation, which accompanies both the normal ageing processes as well as progression to preclinical AD (Villemagne et al., 2013; Hersi et al., 2017; Salthouse, 2019). Overall, both the levels of retrieval-based precuneus activation and fMRI task performance decreased mildly over time in our cohort. Some prior cross-sectional studies reported higher task-related activation in precuneus BOLD signal in cognitively normal older adults compared to younger adults (Miller et al., 2008; Maillet and Rajah, 2014; Soch et al., 2021). This could, however, be related to higher amyloid accumulation or vascular effects that are often not accounted for in studies on normal ageing. Elman and colleagues discussed potential compensatory increases in activation in occipital and parietal areas in amyloid-positive compared to amyloid-negative cognitively normal older adults (Elman et al., 2014). Specific elevated activation might be involved in an attempt at functional compensation to meet task demands (Cabeza et al., 2018), in the presence of early pathological changes and genetic risk. Critically, increased activation could be a compensatory process for a limited time, providing an advantage only in its early phase and leading into a vicious cycle of increasing AD pathology and cognitive decline over time (Jones et al., 2017).

Our results show a relationship between baseline activation, *APOE* genotype, and change in episodic memory performance. Specifically, we observed that the steepest slope of RBANS performance increases over time was present in *APOE4* non-carriers with lower precuneus activity at baseline (compared to all other groups). We note that performance increases probably resemble practice effects, which can even occur when using alternative versions, as in our case (Calamia et al., 2012). Further, participants might show beneficial effects due to their high levels of education (Samson et al., 2023). Our finding suggests that the absence of the *APOE4* allele combined with lower precuneus activity represents a low-risk profile for cognitive decline. With respect to fMRI recognition performance, we observed a decline in fMRI recognition performance in *APOE4* carriers over 48 months that was not present in the *APOE4* non-carriers, with no moderation of cognitive changes by precuneus activity. A stronger decline in recognition memory in cognitively normal *APOE4* carriers compared to non-carriers has been also observed before (Albert et al., 2014; Morrison et al., 2024). It remains open why our findings differ between the different memory measures, with practice effects and moderation by activity for the RBANS memory score but not the fMRI recognition memory task. In summary, our results suggest that *APOE4* carriers show higher risk for memory decline and that this risk might be accentuated in the presence of high precuneus activity.

There are several limitations of our study which should be considered. First, PREVENT-AD is an observational study cohort not initially designed for the testing of our specific hypotheses of whether hyperactivity or activity increases over time predict accumulation of amyloid and tau pathology. However, the study design in this relatively large sample enables us to assess both cross-sectional and longitudinal changes in brain activity via fMRI and related changes in memory performance in a unique manner. Secondly, fMRI is an indirect measure of neural activity and can be influenced by various factors such as the specific task, task demands and vascular contributions (Tsvetanov et al., 2021; Corriveau-Lecavalier et al., 2024). Third, PET data were only available cross-sectionally and subsequent to the fMRI with varying inter-scan intervals. As a result, we cannot comment on amyloid levels concurrent to baseline fMRI, nor how pathology levels may change over time in relation to activity levels. Future analyses could incorporate novel longitudinal plasma markers of amyloid and tau, as recent data showed faster increase in plasma pTau181 levels over time in *APOE4* carriers compared to non-carriers in the PREVENT-AD cohort (Yakoub et al., 2023). This, in addition to assessing the interaction of *APOE4* and intrinsic functional connectivity between PMC regions using longitudinal resting-state fMRI, could highlight other aspects of the complex relationship between functional features and cognitive performance not captured by the current analyses.

In conclusion, our results suggest that greater precuneus activation during memory retrieval is linked to higher subsequent amyloid burden in cognitively normal *APOE4* carriers. Further, the absence of the *APOE4* allele in combination with lower precuneus activation could represent a beneficial low-risk profile for future cognitive decline. These findings could advance ongoing research on pharmacological or non-invasive brain stimulation interventions targeting aberrant activation as a therapeutic target for early AD, which is of significant clinical interest in the context of the emergence of the first disease modifying therapies for amyloid accumulation (Budd Haeberlein et al., 2022; Sims et al., 2023; van Dyck et al., 2023). Our study, therefore, represents a timely exploration into the complex dynamics of precuneus activation, *APOE* genotype, amyloid, and cognition in older adults at risk for AD.

## Acknowledgments

We want to thank the participants of the PREVENT-AD study for their time and effort as well as the researchers involved in building up the cohort https://preventad.loris.ca/acknowledgements/acknowledgements.php?DR=7.0. This work was supported by the German Research Foundation (DFG; Project-ID 425899996, CRC1436 to A.M and E.N.M; Project-ID 362321501, RTG 2413 to A.M. and L.F.).

