## Supplementary Material for "Precuneus activity during retrieval is positively associated with amyloid burden in cognitively normal older *APOE4* carriers"

Conflict of interest statement: The authors declare no competing financial interests.
Acknowledgments: We want to thank the participants of the PREVENT-AD study for their time and effort as well as the researchers involved in building up the cohort https://preventad.loris.ca/acknowledgements/acknowledgements.php?DR=7.0. This work was supported by the German Research Foundation (DFG; Project-ID 425899996, CRC1436 to A.M and E.N.M; Project-ID 362321501, RTG 2413 to A.M. and L.F.).

**Key Words:** Amyloid, *APOE4*, Episodic Memory Retrieval, Functional Hyperactivity, Multimodal Neuroimaging, Precuneus

### Supplementary

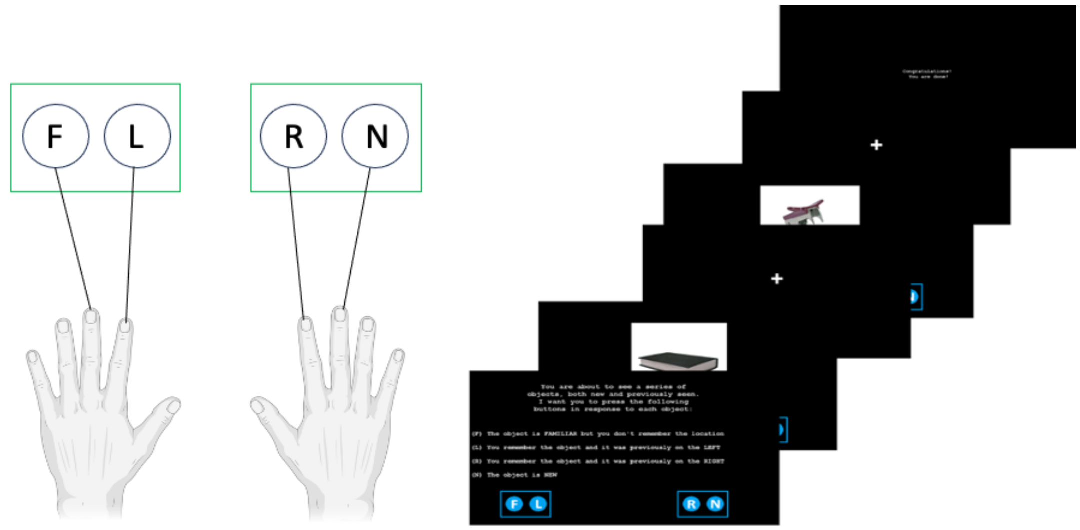


**Supp. Fig. S1. Episodic memory retrieval paradigm:** Following the encoding paradigm, participants were presented with 48 'Old' (previously shown at encoding) and 48 'New' coloured object stimuli presented in the centre of the computer screen. Participants were given a 4-alternative response: "Familiar" (F) to indicate that the object is familiar, but the participant cannot remember whether it was on the left or right, "Remember-Left" (L) to indicate that the participant remembered the object and that it was on the left, "Remember-Right" (R) to indicate that the participant remembered the object and that it was on the right, and "New" (N) to indicate that the object was not previously presented. The presentation rate was 4 seconds.

| | fMRI Precuneus Baseline | | | | | | | |
|---|---|---|---|---|---|---|---|---|
| *Predictors* | *Estimates* | *std. Error* | *std. Beta* | *standardized std. Error* | *CI* | *standardized CI* | *Statistic* | *p* |
| (Intercept) | 1.44 | 3.20 | -0.15 | 0.11 | -4.88 – 7.76 | -0.37 – 0.08 | 0.45 | 0.653 |
| APOE4 Group [carrier] | 0.05 | 0.17 | 0.05 | 0.16 | -0.29 – 0.40 | -0.27 – 0.37 | 0.30 | 0.766 |
| Age Baseline | 0.00 | 0.00 | 0.05 | 0.08 | -0.00 – 0.00 | -0.11 – 0.22 | 0.65 | 0.515 |
| Education Baseline | 0.01 | 0.03 | 0.03 | 0.08 | -0.04 – 0.06 | -0.13 – 0.18 | 0.35 | 0.725 |
| Sex [male] | 0.44 | 0.19 | 0.41 | 0.18 | 0.06 – 0.82 | 0.06 – 0.76 | 2.30 | 0.023 |
| Precuneus GMV | -1.97 | 3.45 | -0.05 | 0.09 | -8.79 – 4.84 | -0.22 – 0.12 | -0.57 | 0.568 |
| Observations | 165 | | | | | | | |
| $R^2$ / $R^2$ adjusted | 0.056 / 0.026 | | | | | | | |

**Supp. Tab. S1. Linear model of effects on baseline activation**: Precuneus activation at baseline was used as dependent variable, APOE4 status, Age at baseline, education, sex and precuneus grey matter volume were used as independent variables. Males showed higher activity at baseline (see also Supp. Fig. S2). GMV = grey matter volume. CI = 95% confidence interval.

| | fMRI Precuneus Slope | | | | | | | | | |
|---|---|---|---|---|---|---|---|---|---|---|
| *Predictors* | *Estimates* | *std. Error* | *std. Beta* | *standardized std. Error* | *CI* | *standardized CI* | *Statistic* | *std. Statistic* | *p* | *std. p* |
| (Intercept) | -0.76 | 2.43 | -0.12 | 0.08 | -5.57 – 4.05 | -0.29 – 0.05 | -0.31 | -1.39 | 0.754 | 0.167 |
| Time | -0.00 | 0.00 | -0.08 | 0.05 | -0.00 – 0.00 | -0.18 – 0.02 | -1.64 | -1.64 | 0.102 | 0.102 |
| Sex [male] | 0.46 | 0.17 | 0.23 | 0.13 | 0.13 – 0.79 | -0.03 – 0.50 | 2.74 | 1.72 | 0.007 | 0.088 |
| APOE4 Group [carrier] | 0.08 | 0.15 | 0.09 | 0.12 | -0.22 – 0.38 | -0.16 – 0.33 | 0.54 | 0.72 | 0.589 | 0.474 |
| Age Baseline | 0.00 | 0.00 | 0.08 | 0.06 | -0.00 – 0.00 | -0.04 – 0.21 | 1.29 | 1.29 | 0.199 | 0.199 |
| Education Baseline | 0.00 | 0.02 | 0.01 | 0.06 | -0.03 – 0.04 | -0.11 – 0.13 | 0.16 | 0.16 | 0.875 | 0.875 |
| Precuneus GMV | 0.49 | 2.63 | 0.01 | 0.06 | -4.71 – 5.68 | -0.11 – 0.14 | 0.18 | 0.18 | 0.854 | 0.854 |
| Time by Sex [male] | -0.00 | 0.00 | -0.20 | 0.08 | -0.00 – -0.00 | -0.35 – -0.05 | -2.61 | -2.61 | 0.009 | 0.009 |
| Time by APOE4 Group [carrier] | 0.00 | 0.00 | 0.01 | 0.08 | -0.00 – 0.00 | -0.14 – 0.17 | 0.19 | 0.19 | 0.852 | 0.852 |
| **Random Effects** | | | | | | | | | | |
| $\sigma^2$ | 0.77 | | | | | | | | | |
| $\tau_{00\ Subject}$ | 0.39 | | | | | | | | | |
| ICC | 0.33 | | | | | | | | | |
| $N_{Subject}$ | 151 | | | | | | | | | |
| Observations | 534 | | | | | | | | | |

**Supp. Tab. S2. Linear model of effects on change in precuneus activation including the interaction term of time by *APOE* genotype and time by sex**: Precuneus activation over time (slope) was used as dependent variable, time, APOE4 status, age at baseline, education, sex, precuneus grey matter volume, and the interaction of time by *APOE* as well as of time by sex were used as independent variables. There was only a significant time interaction with sex as shown in Supp. Fig. S2. GMV = grey matter volume. CI = 95% confidence interval.

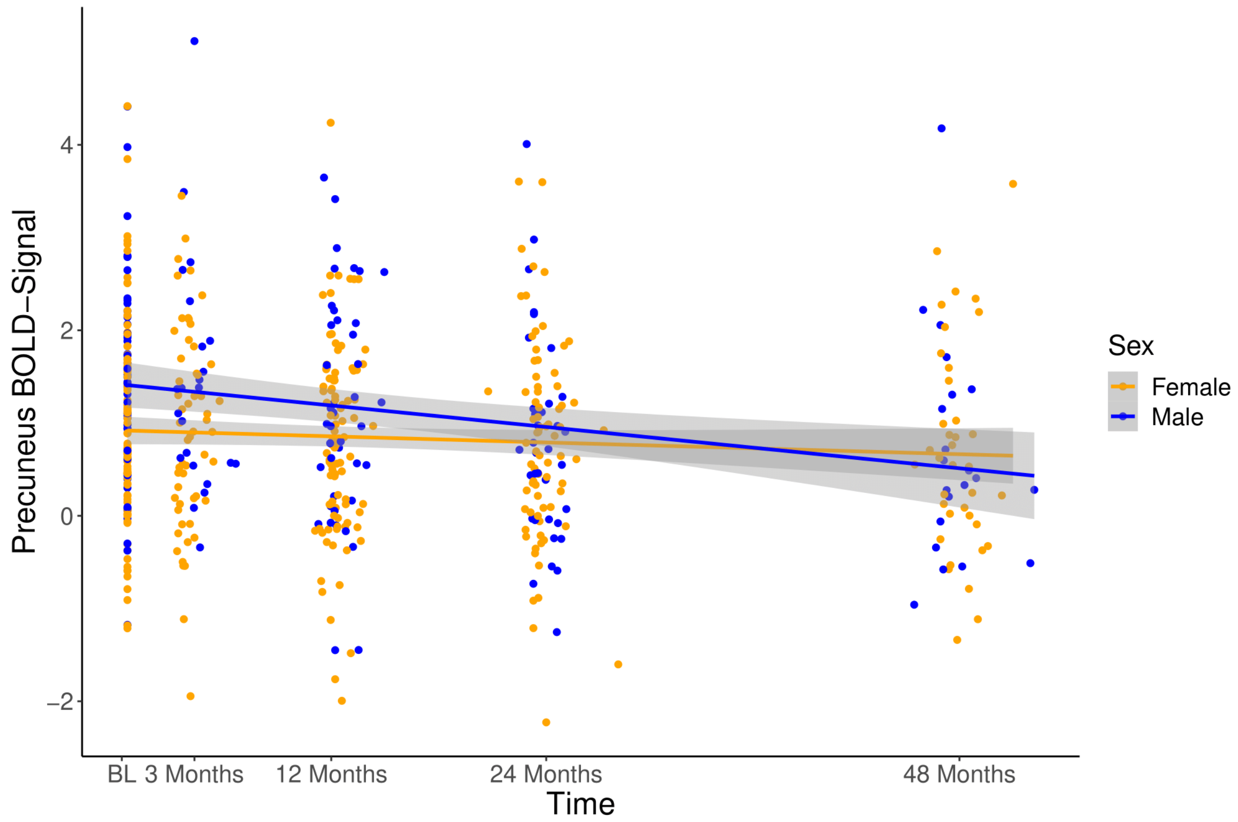


**Supp. Fig. S2. Precuneus retrieval activity at baseline and over time differs depending on sex:** There was a significant effect of sex (β=0.41, [95% 0.06 – 0.76], t=2.297, *p*=0.023), with males showing higher baseline precuneus activation. There was a significant interaction of time by sex (β=-0.20, t=-2.607, *p*=0.009, 95%CI (-0.35, -0.05)), with a decrease in activity over time in males (β=-0.27, [95% -0.39, -0.15], t=-4.591, *p*<0.001; Fig.2B) but no significant change in females (β=-0.08, [95% -0.17, 0.01], t=-1.735, *p*=0.084).

| | Whole brain amyloid PET burden | | | | | | | |
|---|---|---|---|---|---|---|---|---|
| *Predictors* | *Estimates* | *std. Error* | *std. Beta* | *standardized std. Error* | *CI* | *standardized CI* | *Statistic* | *p* |
| (Intercept) | 0.05 | 0.28 | 0.07 | 0.10 | -0.50 – 0.60 | -0.12 – 0.25 | 0.18 | 0.860 |
| fMRI Precuneus Baseline | 0.02 | 0.01 | 0.20 | 0.08 | 0.00 – 0.03 | 0.05 – 0.36 | 2.54 | 0.012 |
| Age Baseline | 0.00 | 0.00 | 0.06 | 0.08 | -0.00 – 0.00 | -0.10 – 0.22 | 0.76 | 0.450 |
| Education Baseline | -0.00 | 0.00 | -0.13 | 0.08 | -0.01 – 0.00 | -0.29 – 0.02 | -1.74 | 0.083 |
| Sex [male] | -0.02 | 0.02 | -0.21 | 0.18 | -0.06 – 0.01 | -0.57 – 0.15 | -1.14 | 0.256 |
| Time Baseline MRI to PET | 0.00 | 0.00 | 0.06 | 0.08 | -0.00 – 0.00 | -0.09 – 0.22 | 0.79 | 0.432 |
| Precuneus GMV | 0.09 | 0.31 | 0.03 | 0.09 | -0.51 – 0.70 | -0.14 – 0.20 | 0.31 | 0.758 |
| Observations | 165 | | | | | | | |
| $R^2$ / $R^2$ adjusted | 0.067 / 0.032 | | | | | | | |

**Supp. Tab. S3**. **Linear model of effects on amyloid PET burden predicted by activation at baseline**: Box-Cox corrected amyloid PET burden was used as dependent variable, fMRI precuneus activation at baseline, age at baseline, education, sex, time between baseline MRI and PET, and grey matter volume were used as independent variables. GMV = grey matter volume. CI = 95% confidence interval.

| | Whole brain amyloid PET burden | | | | | | | |
|---|---|---|---|---|---|---|---|---|
| *Predictors* | *Estimates* | *std. Error* | *std. Beta* | *standardized std. Error* | *CI* | *standardized CI* | *Statistic* | *p* |
| (Intercept) | 0.20 | 0.29 | 0.03 | 0.10 | -0.37 – 0.77 | -0.17 – 0.23 | 0.70 | 0.487 |
| fMRI Precuneus Slope | 1.05 | 0.51 | 0.17 | 0.08 | 0.05 – 2.05 | 0.01 – 0.34 | 2.08 | 0.039 |
| Age Baseline | 0.00 | 0.00 | 0.01 | 0.09 | -0.00 – 0.00 | -0.16 – 0.18 | 0.11 | 0.916 |
| Education Baseline | -0.00 | 0.00 | -0.15 | 0.08 | -0.01 – 0.00 | -0.31 – 0.01 | -1.87 | 0.064 |
| Sex [male] | -0.01 | 0.02 | -0.10 | 0.19 | -0.05 – 0.03 | -0.48 – 0.28 | -0.51 | 0.609 |
| Time Baseline MRI to PET | 0.00 | 0.00 | 0.06 | 0.08 | -0.00 – 0.00 | -0.11 – 0.22 | 0.70 | 0.486 |
| Precuneus GMV | 0.02 | 0.31 | 0.01 | 0.09 | -0.60 – 0.64 | -0.17 – 0.19 | 0.06 | 0.954 |
| Observations | 151 | | | | | | | |
| $R^2$ / $R^2$ adjusted | 0.060 / 0.020 | | | | | | | |

**Supp. Tab. S4**. **Linear model of effects on amyloid PET burden predicted by activation over time**: Box-Cox corrected amyloid PET burden was used as dependent variable, fMRI precuneus activation over time (slope), age at baseline, education, sex, time between baseline MRI and PET, and grey matter volume were used as independent variables. GMV = grey matter volume. CI = 95% confidence interval.

| | Whole brain amyloid PET burden | | | | | | | | | |
|---|---|---|---|---|---|---|---|---|---|---|
| *Predictors* | *Estimates* | *std. Error* | *std. Beta* | *standardized std. Error* | *CI* | *standardized CI* | *Statistic* | *std. Statistic* | *p* | *std. p* |
| (Intercept) | -0.26 | 0.26 | -0.26 | 0.11 | -0.78 – 0.25 | -0.47 – -0.05 | -1.01 | -2.50 | 0.316 | 0.014 |
| fMRI Precuneus Baseline | 0.00 | 0.01 | 0.05 | 0.10 | -0.01 – 0.02 | -0.15 – 0.25 | 0.49 | 0.49 | 0.622 | 0.622 |
| APOE4 Group [carrier] | 0.05 | 0.02 | 0.80 | 0.15 | 0.01 – 0.09 | 0.51 – 1.09 | 2.37 | 5.48 | 0.019 | <0.001 |
| Age Baseline | 0.00 | 0.00 | 0.14 | 0.08 | -0.00 – 0.00 | -0.01 – 0.29 | 1.81 | 1.81 | 0.073 | 0.073 |
| Education Baseline | -0.00 | 0.00 | -0.09 | 0.07 | -0.01 – 0.00 | -0.23 – 0.05 | -1.31 | -1.31 | 0.191 | 0.191 |
| Sex [male] | -0.02 | 0.02 | -0.20 | 0.17 | -0.05 – 0.01 | -0.53 – 0.13 | -1.18 | -1.18 | 0.239 | 0.239 |
| Time Baseline MRI to PET | 0.00 | 0.00 | 0.04 | 0.07 | -0.00 – 0.00 | -0.10 – 0.18 | 0.58 | 0.58 | 0.562 | 0.562 |
| Precuneus GMV | 0.34 | 0.28 | 0.10 | 0.08 | -0.22 – 0.90 | -0.06 – 0.25 | 1.20 | 1.20 | 0.232 | 0.232 |
| fMRI Precuneus Baseline × APOE4 Group [carrier] | 0.03 | 0.01 | 0.29 | 0.14 | 0.00 – 0.05 | 0.00 – 0.57 | 2.00 | 2.00 | 0.047 | 0.047 |
| Observations | 165 | | | | | | | | | |
| $R^2$ / $R^2$ adjusted | 0.235 / 0.196 | | | | | | | | | |

**Supp. Tab. S5**. **Linear model of effects on amyloid PET burden predicted by activation at baseline and *APOE4* group**: Box-Cox corrected amyloid PET burden was used as dependent variable, fMRI precuneus activation at baseline, *APOE4* group, age at baseline, education, sex, time between baseline MRI and PET, grey matter volume, and the interaction of activation at baseline by *APOE4* group were used as independent variables. GMV = grey matter volume. CI = 95% confidence interval.

| | Whole brain amyloid PET burden | | | | | | | |
|---|---|---|---|---|---|---|---|---|
| *Predictors* | *Estimates* | *std. Error* | *std. Beta* | *standardized std. Error* | *CI* | *standardized CI* | *Statistic* | *p* |
| (Intercept) | -0.16 | 0.27 | -0.30 | 0.11 | -0.69 – 0.37 | -0.51 – -0.08 | -0.61 | 0.546 |
| fMRI Precuneus Slope | -0.22 | 0.64 | -0.04 | 0.10 | -1.48 – 1.03 | -0.24 – 0.17 | -0.35 | 0.727 |
| APOE4 Group [carrier] | 0.08 | 0.01 | 0.79 | 0.15 | 0.05 – 0.11 | 0.49 – 1.10 | 5.17 | <0.001 |
| Age Baseline | 0.00 | 0.00 | 0.10 | 0.08 | -0.00 – 0.00 | -0.06 – 0.26 | 1.27 | 0.205 |
| Education Baseline | -0.00 | 0.00 | -0.11 | 0.07 | -0.01 – 0.00 | -0.26 – 0.03 | -1.54 | 0.126 |
| Sex [male] | -0.01 | 0.02 | -0.07 | 0.17 | -0.04 – 0.03 | -0.41 – 0.27 | -0.39 | 0.695 |
| Time Baseline MRI to PET | 0.00 | 0.00 | 0.03 | 0.08 | -0.00 – 0.00 | -0.12 – 0.18 | 0.36 | 0.716 |
| Precuneus GMV | 0.29 | 0.29 | 0.08 | 0.08 | -0.28 – 0.86 | -0.08 – 0.25 | 1.00 | 0.319 |
| fMRI Precuneus Slope by APOE4 Group [carrier] | 2.38 | 0.90 | 0.39 | 0.15 | 0.59 – 4.16 | 0.10 – 0.69 | 2.63 | 0.009 |
| Observations | 151 | | | | | | | |
| $R^2$ / $R^2$ adjusted | 0.238 / 0.196 | | | | | | | |

**Supp. Tab. S6**. **Linear model of effects on amyloid PET burden predicted by activation over time and *APOE4* group**: Box-Cox corrected amyloid PET burden was used as dependent variable, fMRI precuneus activation over time (slope extracted from linear mixed model), *APOE4* group, age at baseline, education, sex, time between baseline MRI and PET, grey matter volume, and the interaction of activation over time (slope) by *APOE4* group were used as independent variables. GMV = grey matter volume. CI = 95% confidence interval.

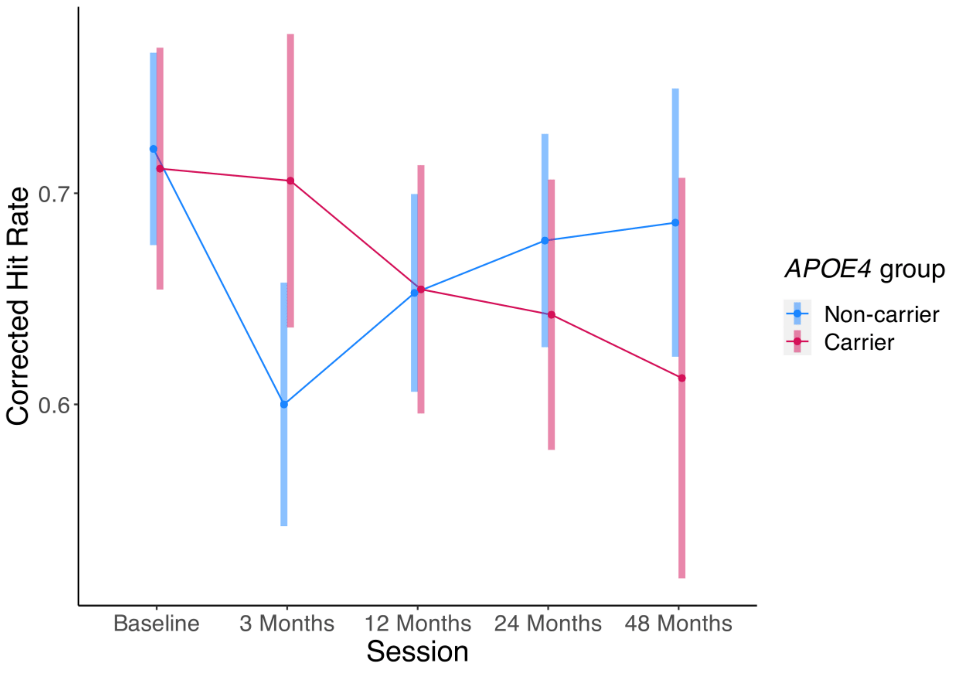


**Supp. Fig. S3. Interaction of *APOE* genotype and session on corrected hit rate.** Results of a linear mixed model show a significant interaction of *APOE* genotype and session on the corrected hit rate, such that *APOE4* non-carriers showed lower performance at the 3-month follow up (t(546)=-2.326, *p*=0.020, SE=0.05, [95% CI -0.20, -0.02]), but here was descriptively higher performance in non-carriers for the 24-month (t(499)=0.856, *p*=0.392, SE=0.04, [95% CI -0.05, 0.12]) and 48-month (t(582)=1.273, *p*=0.204, SE=0.06, [95% CI -0.04, 0.19]) follow-up. Regarding the slopes over time for the corrected hit rate, there was a significant effect of *APOE* group (F(1,148)=89.537, *p*<0.001), with *APOE4* carriers (mean=-0.03, SD=0.02) showing a steeper slope (i.e. decline over time) than non-carriers (mean=-0.01, SD=0.01).

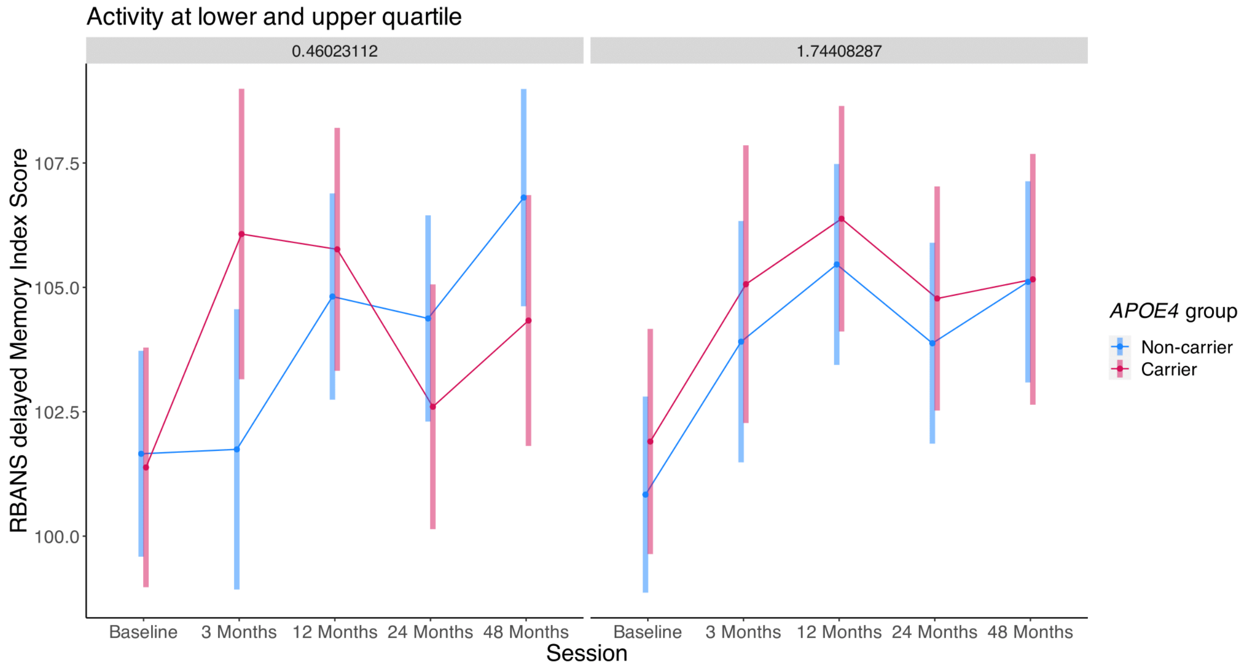


**Supp. Fig. S4. Three-way interaction of *APOE* genotype, precuneus baseline activation and session on RBANS performance.** Results of a linear mixed model show a significant interaction of *APOE* genotype, session, and baseline precuneus retrieval activity on the delayed index memory score. Post-hoc analyses on cross-sectional RBANS performance at follow-up visits between *APOE4* groups and high and low baseline precuneus activation (fixed at the 25% quartile=0.460 and 75% quartile=1.744) revealed an interaction of *APOE4* group and session for lower activation. With lower baseline activation, there is higher performance for *APOE4* carriers compared to non-carriers at the 3-month follow up (t(690)=-2.124, $p$=0.034, SE=2.04, [95% -8.33, -0.33]; not corrected for multiple comparisons), but descriptively higher performance for non-carriers at the 24-month (t(507)=1.109, $p$=0.268, SE=1.60, [95% -1.37, 4.92]) and 48-month (t(544)=1.484, $p$=0.138, SE=1.67, [95% (-0.80, 5.74]) follow ups. There were no differences in performance between *APOE* groups with higher activation (all p>0.05). For N per session, see Fig.1. RBANS = Repeatable Battery for the Assessment of Neuropsychological Status.

| RBANS delayed memory index score | | | | | | | |
|---|---|---|---|---|---|---|---|
| *Session* | *fMRI baseline activation* | *Estimate* | *SE* | *df* | *CI* | *Statistic* | *p* |
| Baseline | Low | 0.28 | 1.58 | 495.70 | -2.83 – 3.38 | 0.17 | 0.863 |
| 3 Months | Low | -4.33 | 2.04 | 689.72 | -8.33 – -0.33 | -2.12 | 0.034 |
| 12 Months | Low | -0.95 | 1.60 | 504.39 | -4.09 – 2.19 | -0.59 | 0.553 |
| 24 Months | Low | 1.78 | 1.60 | 507.18 | -1.37 – 4.92 | 1.11 | 0.268 |
| 48 Months | Low | 2.47 | 1.67 | 543.98 | -0.80 – 5.74 | 1.48 | 0.138 |
| Baseline | High | -1.07 | 1.52 | 501.05 | -4.05 – 1.92 | -0.70 | 0.483 |
| 3 Months | High | -1.16 | 1.88 | 670.54 | -4.85 – 2.53 | -0.62 | 0.539 |
| 12 Months | High | -0.92 | 1.54 | 510.17 | -3.94 – 2.10 | -0.60 | 0.550 |
| 24 Months | High | -0.90 | 1.53 | 507.08 | -3.91 – 2.11 | -0.59 | 0.558 |
| 48 Months | High | -0.05 | 1.64 | 570.74 | -3.27 – 3.17 | -0.03 | 0.975 |
| Observations | | | | | | | |
| $R^2$ / $R^2$ adjusted | | | | | | | |

**Supp. Tab. S7**. **Post-hoc *APOE* group comparisons on marginal means of RBANS performance for low and high baseline precuneus activation**: Post-hoc analyses on cross-sectional RBANS delayed memory index score performance at follow-up visits (sessions) between *APOE4* carriers and non-carriers at low and high baseline precuneus activation (fixed at the 25% quartile=0.460 and 75% quartile=1.744) were conducted after a significant 3-way-interaction of *APOE* genotype, precuneus baseline activation and session on RBANS performance. RBANS = Repeatable Battery for the Assessment of Neuropsychological Status. SE = standard error. CI = 95% confidence interval.
